# Movement Improves the Quality of Temporal Perception and Decision Making

**DOI:** 10.1101/338590

**Authors:** Martin Wiener, Wei Wei Zhou, Wilsaan Joiner

## Abstract

Recent evidence suggest that our experience of time is directly and intrinsically computed within the motor system. Accordingly, a variety of studies have demonstrated that concurrent movement impacts the estimate of temporal duration, such that perceived time is shifted towards the duration of movement. In order to investigate the role of the motor system, we tested human subjects (n=40) on a novel task combining reaching and time perception. In this task, subjects were required to move a robotic manipulandum to one of two physical locations to categorize a concurrently timed suprasecond auditory stimulus as “long” or “short”, relative to a running average criterion. Critically, subjects were divided into two groups: one in which movement during the interval was unrestricted and subjects could move freely to their choice, and one in which they were restrained from moving until the stimulus interval had elapsed. Our results revealed a higher degree of precision for subjects in the free-moving group. By further decomposing choice and response time data with a drift diffusion model of decision making, we identified the source of this change to a shift in the response threshold for free-moving subjects. Analysis of movement parameters revealed that eventual choice could be determined by movement parameters (e.g. trajectory, force) before the response was made, and even before the interval had elapsed, and that the variability of these movements was lower in the unrestrained group. A further experiment (n=14) verified that these findings were not due to proximity to the target, counting strategies, bias, or movement length. Our findings suggest that perceptual timing may be instantiated within the motor system as an ongoing readout of timing judgment and confidence.

## Introduction

A critical aspect of behavior is that mobile organisms must be able to precisely determine *where* and *when* to move (Rosenbaum, 2009). A better understanding of the mechanisms underlying precise movement timing and action planning is therefore crucial to understanding how we interact with the world around us. A majority of studies focusing on action planning and reaching movements have been focused over the past thirty years on spatial aspects of reaching (Wolpert et al., 2011). Specifically, there is interest regarding the locations where subjects will move to, and how subjects will update their movements to particular locations when the targets they are reaching for are either unknown or changed. These studies further focus on motor adaptation paradigms, in which the parameters of a movement trajectory are altered, unknown to the subject, such that a subject does not arrive at the location they expected to, and so must update their internal model of movement in order to accommodate these changes (Wei and Körding, 2009; Gallivan et al., 2017; Hosseini et al., 2017; Sing et al., 2009). The results of these studies provide important insights into how the motor system both adapts and accommodates changes in movement trajectories, and have provided dissociations between the preparation, planning, and executions of movements (Shadmehr et al., 2010; Wong et al., 2016; Haith et al., 2016).

More recently, movement-related paradigms have been adopted to examine perceptual decision making (Freeman et al., 2011). Here, subjects are required to move a robotic manipulandum to one of two locations, each of which corresponds to a particular choice (Barca and Pezzulo, 2012; Finkbeiner and Heathcote, 2016; Hagura et al., 2017). By exploiting the dynamics of reaching trajectories (e.g., the velocity and force during the movement) different parameters can be measured to reveal a moment-by-moment sampling of the motor system and the deliberation process (Wolpert and Landy, 2012). For example, using this paradigm researchers have shown that when a stimulus is ambiguous, subjects may initially reach for a particular target, and then deviate from this path towards the opposite choice (Resulaj et al., 2009). Such “changes of mind” have been explained by postulating a post-decisional component in sequential sampling, where subjects continue to accumulate evidence in favor of competing options.

A critical aspect of the above studies is the timing of motor movements. Indeed, timing is inexorably tied to movement, as for any movement to take place, temporal precision is needed to coordinate between different limb movements and the environment (Todorov, 2004). Remarkably, recent studies show that our perception of time depends on whether we are moving or not (Imaizumi and Asai, 2017) (Tomassini et al., 2012)(Tomassini et al., 2014) (Yokosaka et al., 2015) (Press et al., 2014) (Yon et al., 2017)(Hagura et al., 2012)(Tomassini and Morrone, 2016). Prolonged movements serve to expand the perceived duration of a concurrently presented stimulus (Yokosaka et al., 2015), and perceptual judgments of time tend to gravitate towards the length of our motor movements (Press et al., 2014), even when these movements are not tied to the task in any way (Yon et al., 2017). These distortions can also occur when viewing the movements of other people, and can further be altered by the sense of agency a viewer has during the observation (Imaizumi and Asai, 2017). Further, even the preparation of a movement is sufficient to induce temporal distortions, such that stimuli are perceived as longer when we are preparing to move than when not (Hagura et al., 2012). These distortions are also dependent on both the direction and speed of movement being planned or executed, with faster movements and those away from the body leading to longer perceived durations, and slower movements and those towards the body leading to shorter perceived durations (Yokosaka et al., 2015; Tomassini and Morrone, 2016). Finally, movement distortions affect concurrently perceived stimulus durations regardless of the stimulus modality in which they are presented (Tomassini et al., 2014, 2012; Tomassini and Morrone, 2016), suggesting that motor movements serve a supramodal role in the processing of temporal duration.

While the above findings suggest that movements can serve to distort perceptual timing, there is also evidence to suggest that it can enhance it. Indeed, evidence has shown that, during movement preparation, in addition to an expanded perception of time, there is also an increase in temporal fidelity, such that subjects can more easily perceive visual flicker and rapidly presented stimuli (Hagura et al., 2012). Similarly, when time perception for a particular modality has been altered, such as via adaptation, concurrent movements can serve to re-orient subjects to the appropriate duration (Tomassini et al., 2012). These findings suggest that, when concurrent movement is engaged or simulated, the timing of environmental stimuli adopts computations of the motor system. However, whether this adoption is due to timing being overwritten by the motor system, or enhanced by it – because the motor system itself is the one doing the timing – remains unknown.

In order to determine the involvement of the motor system in temporal perception and decision making, we tested human participants on a reaching paradigm in which they were required to classify auditory stimuli of different lengths into distinct duration categories (Figure 1). Crucially, we manipulated the motor requirements of the task in different groups of subjects, with some being allowed to freely move during stimulus presentation and others being restrained until the stimulus had fully elapsed. The goals of this study were to examine whether (1) concurrent movement during perceptual judgments of time can enhance temporal perception, (2) this enhancement can be tied to specific components of timing, and (3) is linked to intrinsic computations of the motor system.

**Figure 1: Figure 1.**
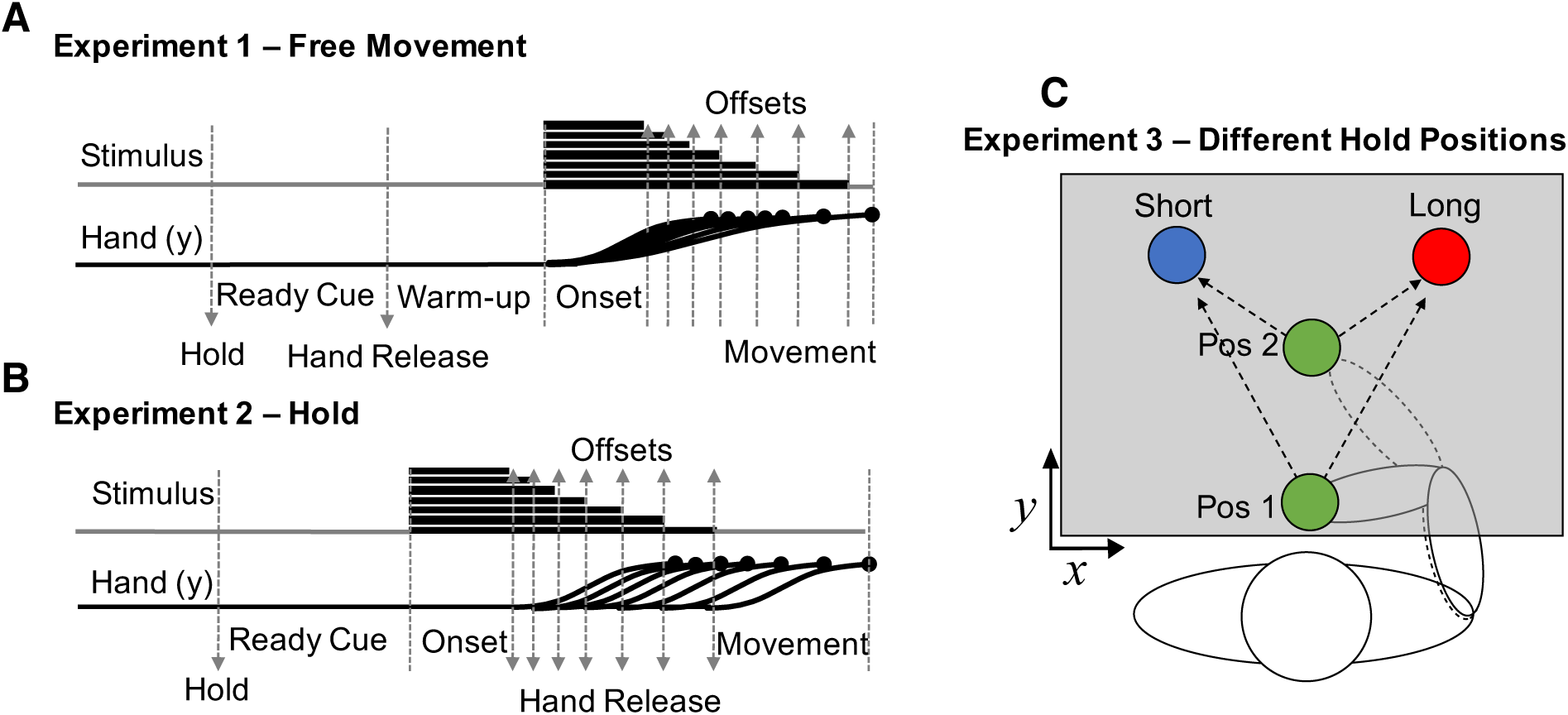
Task design for the three experimental types. In each experiment subjects held the handle of a robotic manipulandum and moved a screen cursor to one of two targets to classify a tone as short (left target) or long (long target). **(A)** In experiment 1 (free-movement), subjects were held in place in the starting position, while a ready cue was presented. Following this, the arm was released for a “warm-up” period, in which subjects could freely move. Onset of the tone proceeded for one of seven possible durations (log-spaced), and subjects were free to move to one of two target locations, but could only enter a location after tone offset occurred. **(B)** In experiment 2 (hold) no warm-up period was provided, and subjects were only released once the tone duration was complete. **(C)** In experiment 3 (different hold positions) subjects performed the hold experiment, but with one of two possible starting positions, in different blocks.

## Results

Our analysis began by measuring psychometric and chronometric functions, in which we measured the proportion of trials in which subjects classified each duration as “long”, and the reaction time for each duration, respectively. Psychometric data were additionally fit with a cumulative gumbel distribution, using the bias-corrected bootstrap method outlined by Wichmann and Hill (2001) and implemented in the *psignifit* toolbox (˙). From this curve, the bisection point (BP) was determined as the duration corresponding to a 0.5 probability of classifying the duration as long. Additionally, the coefficient of variation (CV) was determined as the difference between the 0.75 and 0.25 probability durations (difference limen) divided by the BP. Our initial analysis of these data revealed that subjects in the free movement group were significantly less variable than subjects in the hold group (Figure 2), as characterized by a lower CV [Mann-Whitney U = 124, *p* = 0.012, Cohen’s *d* = 0.69, 10,000 permutations], with no change in the BP [Mann-Whitney U = 196.5, *p* = 0.925] (Figure 4). Additionally, RTs were significantly lower in the free movement group than in the hold group [duration by group interaction: *F*(6,228)=15.743, p<0.001, *η*^2^_p_= 0.293] which had been expected, given that these subjects were allowed to move closer to the targets during tone presentation; yet, unexpectedly, the RT advantage only held for durations longer than the BP, (> 2000 ms), with comparable RTs for durations classified as short. These findings suggests that the allowance of movement during timing conveys a perceptual advantage in temporal bisection.

**Figure 2: Figure 2.**
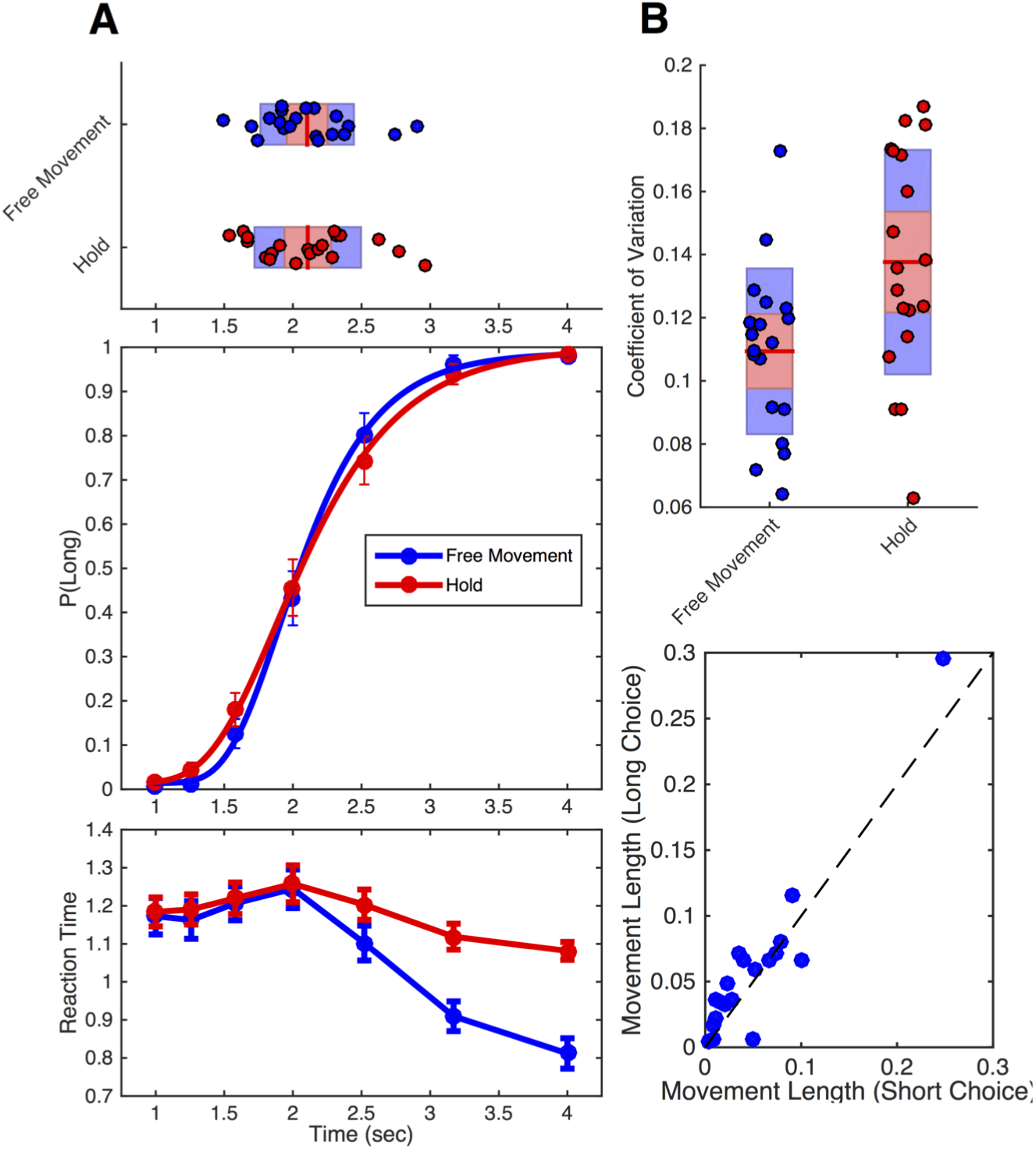
Behavioral data from free-movement and hold experiments. **(A)** The middle panel displays the average proportion of trials on which subjects classified durations as “long” as a function of tested duration. Psychometric curves represent fits to the average data. Red symbols represent data from experiment 2 (hold), blue symbols represent data from experiment 1 (free-movement). The bottom panel displays average chronometric data, with reaction time as a function of duration. Reaction time was notably similar for both groups for durations under two seconds, but then became increasingly faster for free movement subjects. The top panel displays individual bisection points, derived from fitted curves. Boxplots display the mean (red line), bounded by the 95% confidence interval (red shaded region) and the standard deviation (blue region). **(B)** Top panel, coefficient of variation data for both groups, demonstrating significantly lower variability for the free-movement experiment than the hold experiment. Bottom panel, the average movement length for subjects in the free-movement experiment for two second trials classified as “long” or “short”; subjects chose “long” more often when they moved more during the two second tone interval. Results remained significant with removal of the outlier. Error bars represent SE.

**Figure 3: Figure 3.**
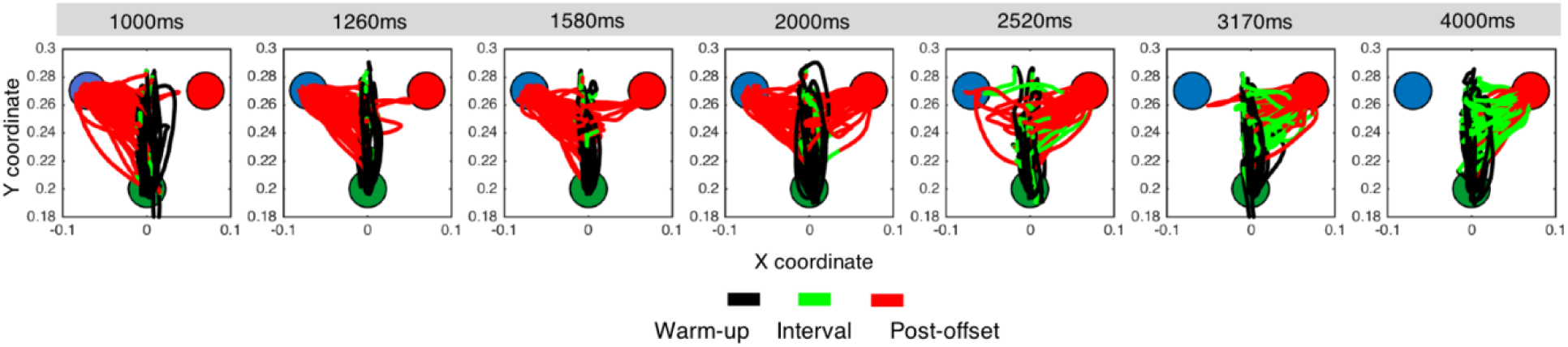
Movement trajectories for an example subject in experiment 1 (free-movement). Layout of the start position (green filled circle), and short (blue filled circle) and long targets (red filled circle) is the same as in Figure 1C. Distance is in meters. Each panel represents the total number of movements for one of seven intervals used in our temporal bisection task. Color coding of traces reflects hand movement through the progression of the given trial (black: warm-up period, green: during the tone duration, red: after the tone until the decision). As can be observed, for relatively “short” durations (< 2000 ms), the majority of movements smoothly progress from the starting position at the bottom of the panel to the short target location (left target). As durations increase (1580, 2000 and 2520 ms), choices split between the short and long locations (note the directions of the red traces). Notably, for relatively “long” durations (> 2000 ms), subject arm movements initially move towards the short location, but then shift towards the long location (right target) once the interval has elapsed past a certain point.

**Figure 4: Figure 4.**
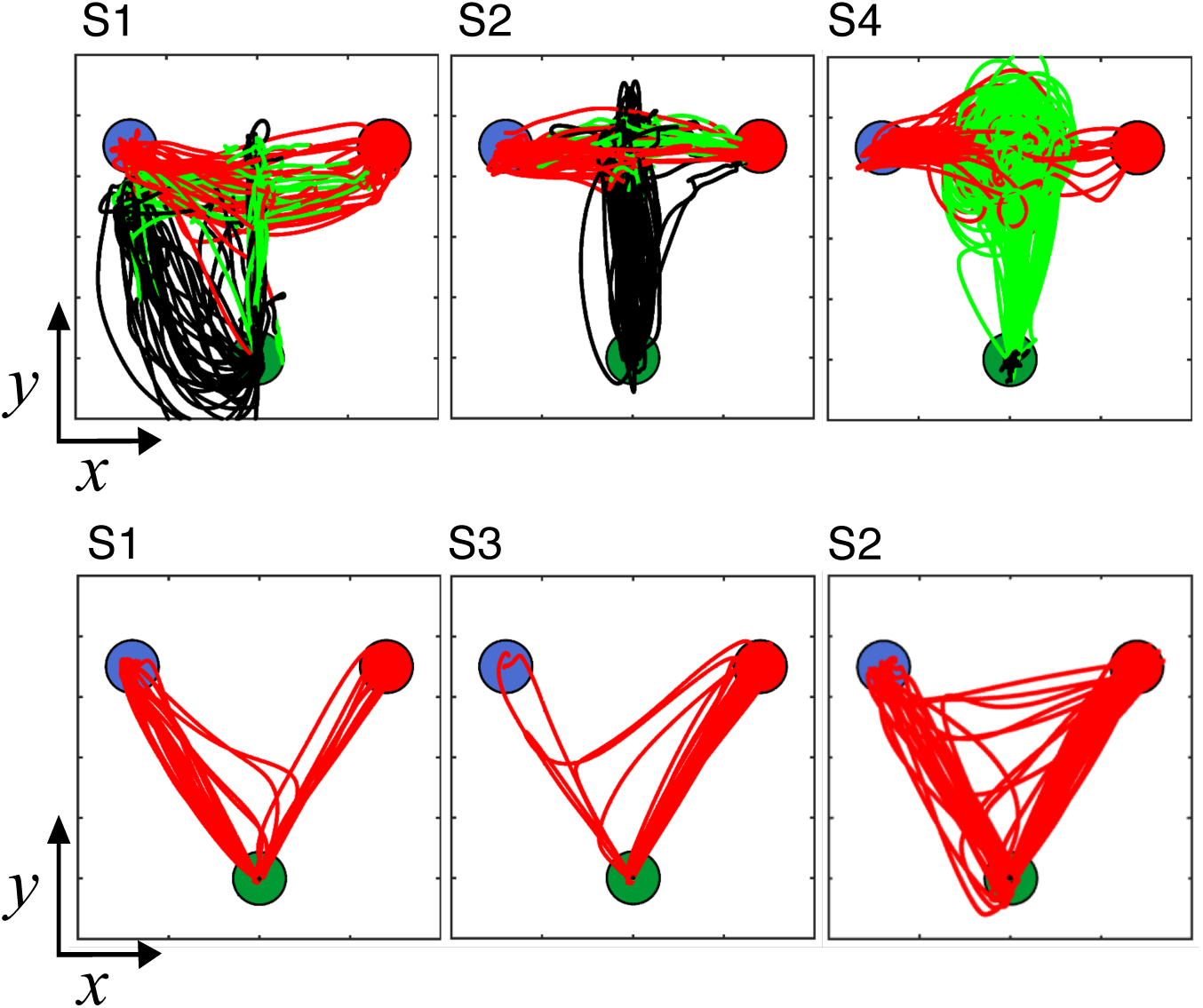
Movement trajectories are idiosyncratic between subjects, yet consistent within subject. Layout of the start position (green filled circle), and short (blue filled circle) and long targets (red filled circle) is the same as in Figure 1C. Color coding of traces reflects hand movement through the progression of the given trial (black: warm-up period, green: during the tone duration, red: after the tone until the decision). (**Top row**) Trajectory data for 2000 ms trials for three example subjects from experiment 1 (free-movement). Classification of a “short” duration is the left target, a “long” classification is the right target. Each subject employed a separate strategy, with one subject (S4) moving in a circular pattern between short and long locations, another (S1) rotating in a leftward arc before moving in between both targets, and a third (S2) moving in an up-down fashion before shifting from the middle to a target location. (**Bottom row**) Three example subjects for 2000 ms trials for three example subjects from experiment 2 (hold). Changes-of-mind are evident, including a shift from long to short (S1), short to long (S3), and both (S2).

Why might subjects perform better during free movement than the hold condition? Examination of the movement trajectories revealed a complex picture into the deliberation process. As expected for the ideal strategy, some subjects moved the cursor closer to the short duration target initially, but then gradually moved closer to the long duration target as the interval on a given trial elapsed (Figure 3). However, across subjects, examination of movement trajectories revealed that nearly every subject adopted a slightly different strategy (Figure 4). Some subjects adopted a “circular” strategy, where they spun the arm in a circle during the warmup and duration presentation phases (Figure 4, S4); others chose a “middle” strategy, keeping the cursor in between the short and long locations and hedging closer to one location or the other (Figure 4, S1). Still others chose an “up-down” strategy, where they moved the cursor along the midline up and down between the two targets before moving to a target at offset (Figure 4, S2). Thus, subject trajectories were idiosyncratic, yet *consistent* within each subject. Given the patterned nature of performance, where some free-movement subjects moved the cursor in a rhythmic-like fashion, one possibility is that subjects were using the arm to adopt a counting strategy. As counting can lead to reductions in the CV (Hinton and Rao, 2004), this may have accounted for the improvement in performance. We analyzed this possibility by collecting the average path length traveled for subjects across trials during each duration. As such, subjects who moved in a rhythmic fashion would have a longer path length than subjects who kept the cursor in a relatively stable location or chose the ideal strategy. These subjects should also have lower CVs than subjects with no patterned movement. Although a negative relationship was observed, no significant correlation between mean path length and CV was found [Spearman rho = -0.184, *p* = 0.446, 10,000 permutations], suggesting that the improvement in CV was not due to a counting/rhythmic strategy. Additionally, no between-subject correlation was found between average movement length and the BP [rho = 0.279], indicating that movement length did not overall shift bias in responses. However, while no effect between subjects was found on the CV, there is the possibility that movement length did still influence choice within subject. Here, we extracted mean movement length for trials classified as “short” vs “long”, only for the middle duration of the stimulus set (2000 ms), at the approximate location of the BP and the most ambiguous point. We observed an impact of movement length, with longer movement lengths when subjects chose “long” than when they chose “short” [Wilcoxon Signed Rank Test, *Z* = -2.173, *p* = 0.027, Cohen’s *d* = 0.753, 10,000 permutations] (Figure 2B botton panel), consistent with previous reports that longer movements are associated with longer perceived intervals (Yokosaka et al., 2015).

For the hold group, although movement trajectories were constrained to ballistic movements to the target at interval offset, notable inter-subject differences were observed (Figure 4, bottom row). The most salient feature here was that these subjects exhibited the same “change-of-mind” pattern as observed in other recent reach-to-touch decision making tasks (van et al., 2016; Resulaj et al., 2009). Specifically, while the majority of trials were characterized by direct movements to a target, an observable subset of trials contained trajectories that began towards one target, and then switched in direction to a different target. These changes-of-mind occurred in both directions (short-to-long; long-to-short), and notably occurred more often for some subjects than others (Figure 6). Previous research has suggested that these changes occur as a re-evaluation of the given evidence after a stimulus has been presented, and further show that, when a subject changes their mind, they often change it to the correct answer Resulaj et al. (2009). In the temporal bisection task, a “correct” choice is relative to the criterion used for classification, which may vary by subject. Yet, the observation of changes-of-mind alone does not explain why these subjects are more variable.

**Figure 5: Figure 5.**
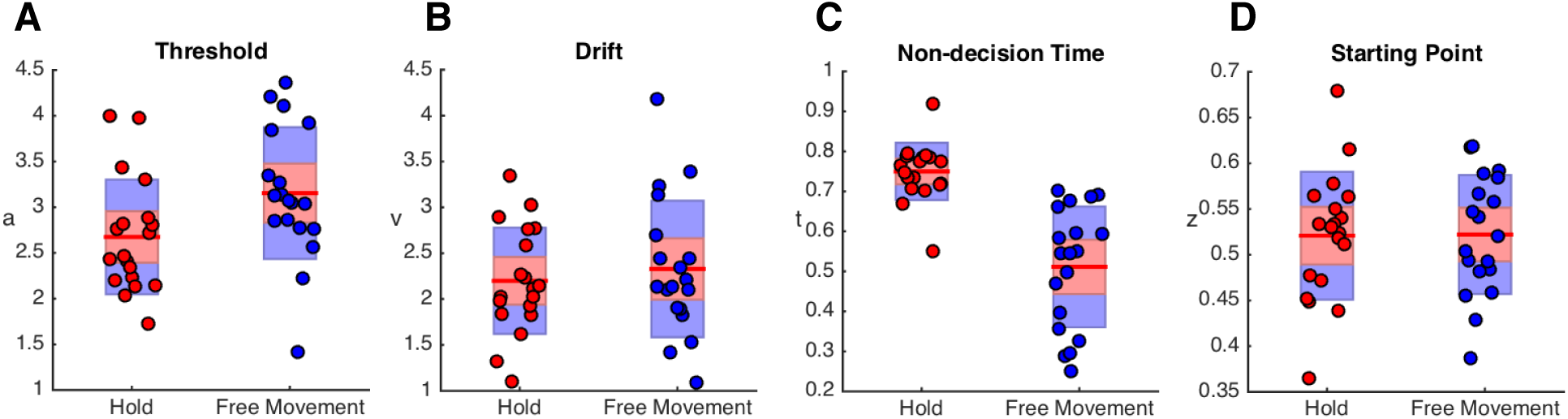
Drift diffusion model results. Presented are the **(A)** threshold, **(B)** drift, **(C)** non-decision time and **(D)** starting point parameters from the drift diffusion modelling of the behavioral results. Red symbols represent individual subjects in experiment 2 (hold), blue symbols represent experiment 1 (free-movement). Boxplots display the mean (red line), bounded by the 95% confidence interval (red shaded region) and the standard deviation (blue region). There was a significant difference between groups for the threshold parameter (*a*) and the non-decision time parameter (*t*).

**Figure 6: Figure 6.**
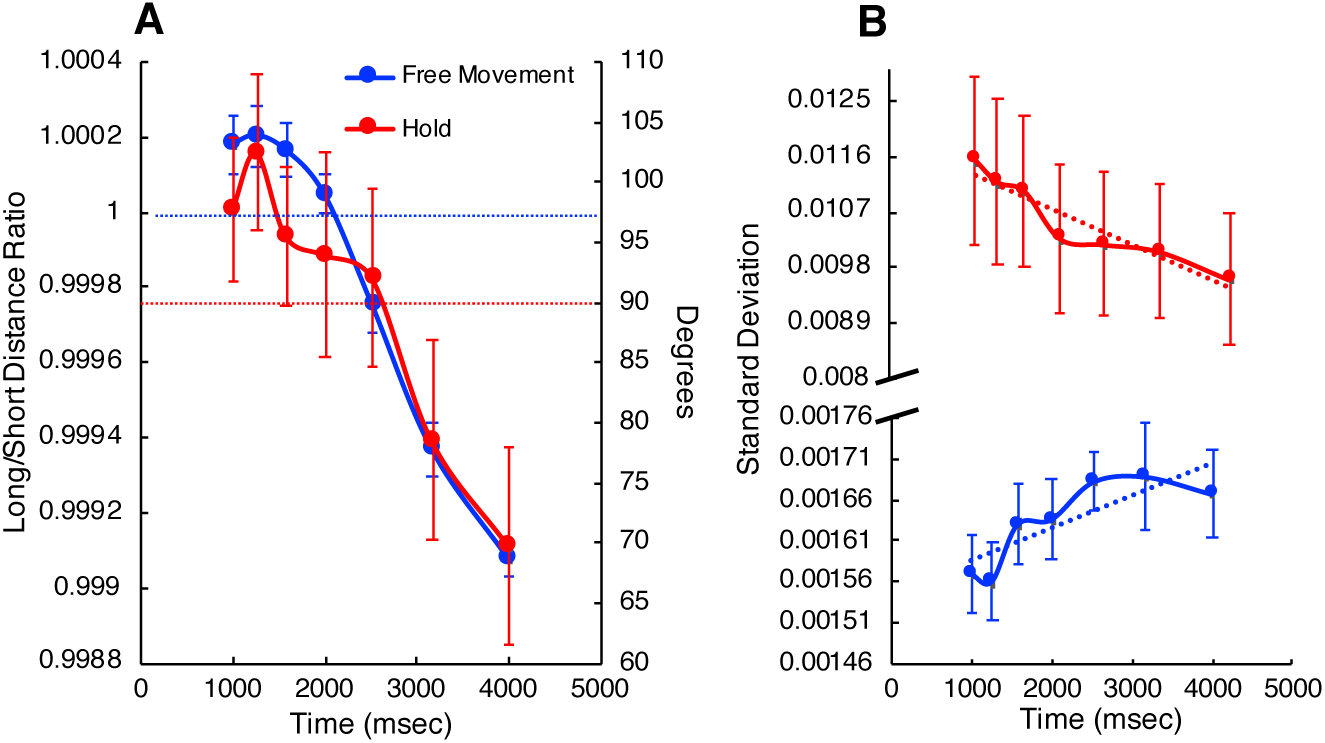
The decision process is represented within the movement trajectories and force patterns. **(A)** Position and force data for both experiments as a function of tone duration. Red symbols represent data from experiment 2 (hold), blue symbols represent data from experiment 1 (free-movement). The left axis displays the ratio of the Euclidean distance from arm position to the long and short targets for the free-movement group; values above 1 (horizontal blue dashed line) indicate the subject was closer to the short location, and vice versa. The right axis displays the force direction, in degrees, that subjects were exerting on the arm at tone offset. Values above 90^<u>o</u>^ (horizontal red dashed line) indicate the subject was pushing towards the short target (105^<u>o</u>^), whereas below 90^<u>o</u>^ indicate pushing towards the long target (75^<u>o</u>^). **(B)** Inter-trial variability (standard deviation) of the arm position was measured for the final 100ms of arm movements across trials for each duration. Free-movement subjects (blue trace) demonstrated linearly increasing SD with tone duration, whereas hold subjects (red trace) displayed linearly decreasing SD. Note the axis break and scale difference in the y-axis, demonstrating that hold subjects exhibited greater variability overall compared to free movement subjects. Error bars represent SE

### Drift Diffusion Modeling

In order to further explain why free-movement subjects may have improved precision over hold subjects, we decomposed choice and RT data with a drift diffusion model of timing and decision making (2ı); (Wiener et al., 2018b). According to this model, subjects accumulate noisy temporal information while a to-be-timed stimulus is active. Once the stimulus has extinguished, subjects then accumulate evidence towards one of two decision boundaries where a response is initiated. Previous studies have demonstrated four features of this model in temporal bisection studies: 1) the separation between the boundaries (*a*) becomes wider with more certainty about the decision; that is, if the interval is very short or very long, more evidence is needed to commit to a decision, whereas if the interval is ambiguous, the level of evidence for deciding short or long is low. 2) The drift-rate (*v*) similarly increases with greater certainty, with faster accumulation of evidence for very short or very long durations, but slower rates for ambiguous durations. 3) The starting point of the decision process (*z*) starts closer to the short duration boundary, but progressively moves closer to the long duration boundary with longer perceived durations. 4) The residual non-decision time (*t*) decreases linearly with longer durations.

To apply our model-based approach, we included routines provided by the HDDM package for Python (Wiecki et al., 2013). As described above, we constructed our model to obtain four measures: threshold (*a*), drift-rate (*v*), non-decision-time (*t*), and starting point (*z*). Each parameter was set to vary by the duration of the presented stimulus. Model fits were obtained via a Maximum Likelihood procedure which minimized the difference between model output and subject choice and reaction time data. The results of our analysis replicated the predicted patterns for all four parameters as described in previous reports (2ı); (Wiener et al., 2018a). Notably, no interaction between experimental group or duration was found for any parameter (all *p* > 0.05); as such, we collapsed across duration for additional comparisons between groups. The results of this analysis revealed a significant effect only on the threshold parameter (*a*) [Mann-Whitney U = 94, *p* = 0.004, *d* = 1.017], as well as the non-decision time parameter (*t*) [Mann-Whitney U = 13, *p* < 0.001, *d* = 2.665] (Figure 5). Specifically, subjects in the free-movement group were characterized by a higher threshold than subjects in the hold group, regardless of duration. No changes were found for the drift or starting point parameters. This finding suggests that the effect of movement was on the decision-stage, rather than on the actual perception of duration (Hagura et al., 2017). Increases in threshold parameters in drift diffusion models are often interpreted as an increase in response caution, where a subject requires more evidence before committing to a particular choice (Voss et al., 2004). In this case, one explanation is that subjects in the free-movement group were afforded more caution in their choice because the freedom of movement afforded them more flexibility to make a decision than subjects in the hold group. Yet, the results do not address if this improvement was due to being physically closer to the response targets, or to the act of moving itself, a point we return to in Experiment 3 described below.

### Movement Dynamics Inform Temporal Estimates

In addition to the above findings, our results suggested that the subject’s choice could be determined by the movement dynamics (Figure 8). To determine this, we first examined the location of the cursor at the offset of the duration signal in experiment 1 (free-movement). For each trial, we calculated the Euclidian distance of the cursor to the short and long targets, and then took their ratio (short distance / long distance); values above 1 indicate the cursor was closer to the short target. Here, we found that, regardless of strategy used, free-movement subjects were closer to the short-duration target at the shortest interval offset, and then gradually shifted towards the long-duration target with longer intervals [*F*(6,114) = 58.912, *p* < 0.001, *η*^2^_p_=0.756]. As expected, this finding suggests that subjects decided which choice they will make before the target duration had elapsed. For the hold group, the position at offset would provide no information, as subjects were held in place until this point in time. Instead, we measured the force being applied to the robotic manipulandum arm handle by the subject. On each trial, we calculated the force being applied in x and y coordinates, and the resultant direction subjects were pushing in. Here, we found that subjects were pushing in the direction of the short target location (105°) for the shortest duration, and then gradually shifted their direction towards the long target location (75°) with increasing duration [*F*(6,114) = 10.720, *p* < 0.001, *η*^2^_p_=0.361]. Both effects were linear, and showed a remarkable degree of correspondence between experimental groups (Figure 6B). A noteworthy feature of this finding is that it indicates that the choice was not simply binary; that is, subjects did not adjust force in a stepwise fashion, as would be expected if motor output simply reflected the crossing of an internal categorical boundary. In other words, *motor output did not merely represent the endpoint of the decision*. Instead, subjects moved along a gradient between the two target locations. This gradient provides a readout of elapsed time from the subject’s movement, and also further highlights the uncertainty subjects had in classifying temporal durations, as durations near the middle of the stimulus set were also near the respective middle locations. Further, it suggests the motor system is continuously evaluating the length of an interval, and can be used as a measurement of subjective internal timekeeping.

**Figure 7: Figure 7.**
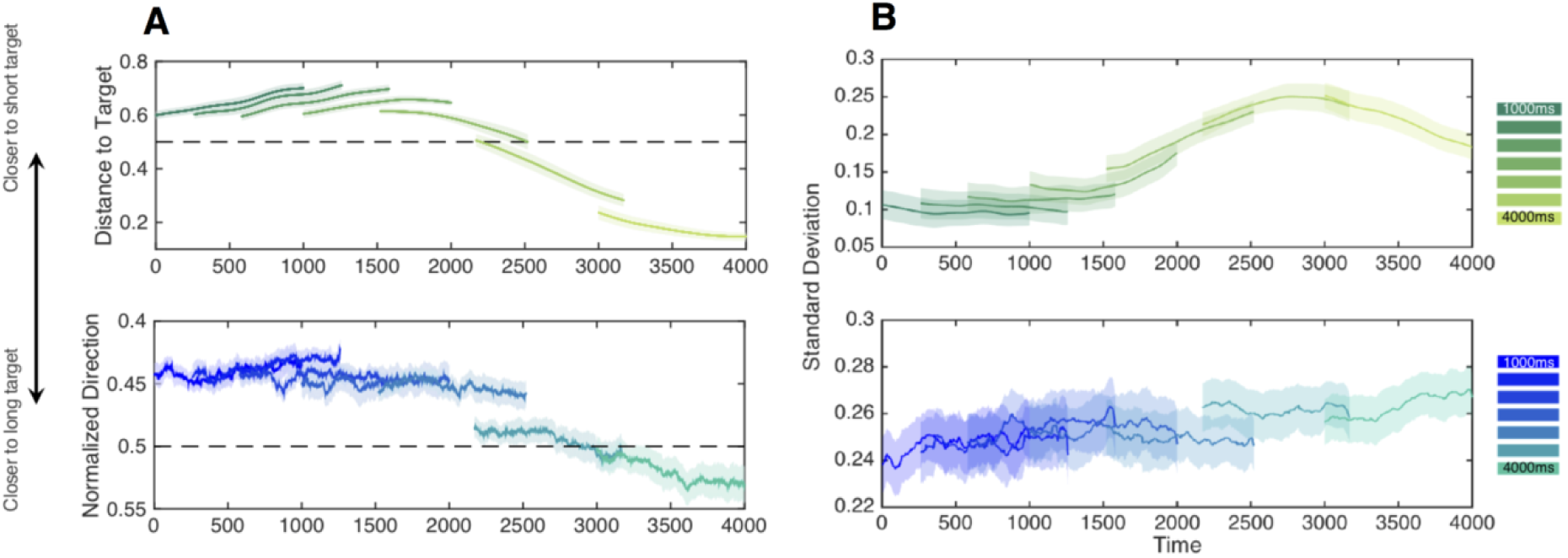
Differences in movement dynamics across tasks. **(A)** Movement dynamics for the free-movement group (top panel) and hold group (bottom panel) during the presentation of each interval. Data are shown for the last 1000ms of each presented interval. Dashed lines represent the 50% inflection point between the long and short duration targets, with values above this line representing a closer relative location (top panel) or a closer force direction (bottom panel) to the short target location. All displayed data are sigmoid transformed. For both sets, movement dynamics display a transition from proximity to the short duration target to the long duration target as the interval elapses; additionally, differences in position or force could be distinguished well before the interval offset. **(B)** Inter-trial variability of movement dynamics displayed in (A). Both panels display the average within-subject standard deviation of movement data from their corresponding left panels. For both groups, inter-trial variability increases as the presented interval elapses, but with different profiles. Further, variability is dramatically lower for the free-movement group as compared to the hold group (note the difference in the vertical scale). Shaded regions display SE.

**Figure 8: Figure 8.**
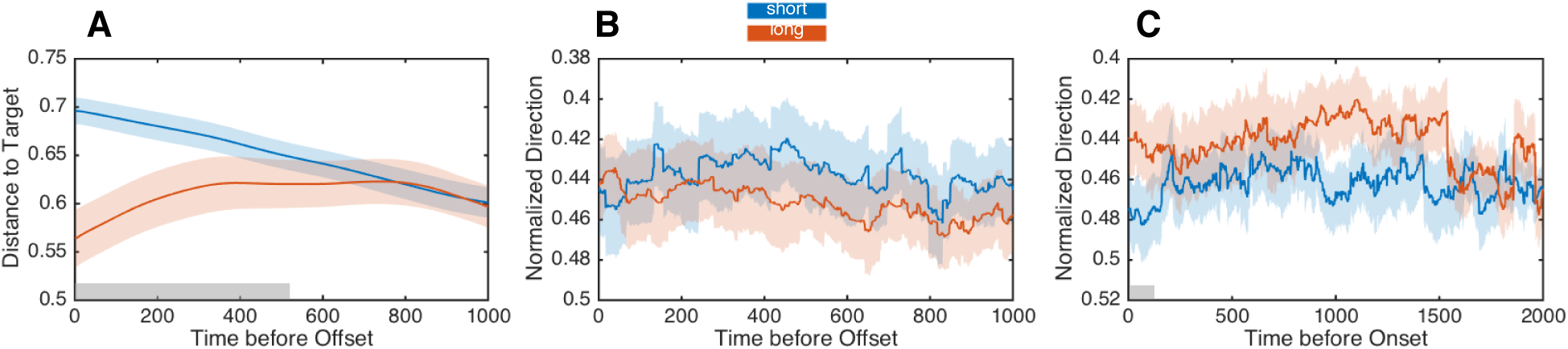
Movement dynamics predict eventual choice. All three panels display data for the middle interval of the stimulus set (2000 ms) **(A)** Normalized movement ratios for the free-movement group. Eventual choice could be determined approximately 500 ms prior to interval offset. **(B)** Normalized force direction for the hold group. In contrast to the free-movement group, no distinguishable difference in direction was observed between short and long response choices. **(C)** Normalized force direction for the hold group during the interval prior to stimulus onset. Here, a distinguishable difference was detected in the final 100ms prior to target onset; note that the direction subjects are exerting force is pointing in the opposite direction of the choice they will eventually make on that trial. Gray shaded regions indicate significance at *p*<0.05 for logistic regression.

Consistent with the above explanation, we examined the variability of movement trajectories as subjects were approaching their chosen target. We measured the inter-trial variability of movement trajectories in the last 100ms of each trial’s movement (Figure 6B). Here, we again found a remarkable difference between groups. For the free-movement group, the average movement variability increased linearly with increasing durations [*F*(6,114) = 2.287, *p* = 0.04, *η*^2^_p_=0.207], such that subjects were more variable in their movements when committing to a longer duration choice. Yet, for the hold group, we found the exact opposite pattern; subjects became *less* variable in their movements with longer durations [*F*(6,114) = 4.638, *p* < 0.001, *η*^2^_p_=0.196]. Additionally, these subjects were dramatically more variable overall in their movement trajectories than in the free-movement group, despite the fact that these measurements were collected from the last 100ms of movement in both groups. One reason for this difference may have been the distance subjects must travel when making a choice. Indeed, movement trajectories are commonly more variable in reaching for further targets (Messier and Kalaska, 1999). Yet, the different relationships between variability with increasing duration cannot alone be explained by this. As subjects were also faster at making reaching movements after longer durations, we suggest a distance/speed tradeoff in variability is the source of this effect. That is, if a target is far away, progressively faster (and more certain) movements become less variable, whereas if a target is close, faster movements are more variable. This finding follows kinematic theories of movement (Plamondon and Alimi, 1997), which suggest a nonlinear effect of the speed/accuracy tradeoff for different movement distances (Hancock and Newell 1985).

To further examine the effects of force and distance ratios in our two experimental groups, we performed a logistic regression analysis of single trial responses with the movement parameters described above. Specifically, we constructed a generalized linear mixed effects model in which the response choice (“short” or “long”) could be predicted by the long/short distance ratio for the free-movement group, or the force direction data for the hold group. We further repeated this analysis in 1ms steps for the 1000ms preceding the offset of the interval for both groups. This time value (1000ms) reflects the largest amount of time from interval offset that distance/force data was available for all intervals in our stimulus set. The first finding of this analysis was that choices could be accurately decoded for the free-movement group for up to 810ms prior to the offset of the interval (all p<0.05, Bonferonni corrected). This finding extended our analysis of the offset points alone, with the movement trajectories demonstrating a good separation between distance ratios well before offset of the interval (Figure 7A). For the hold group, force data were notably noisier; significant periods where force direction predicted choice were detected, but not for any consistent period of time, and only at an uncorrected level (p<0.05).

The above data further suggests that the momentary position or force exertion throughout the interval can serve as an index of the internal timekeeping process. To that end, we also examined inter-trial variability in both measures. If movement dynamics provide a measure of temporal estimation, then the the variability of those measurements should index the variability, and uncertainty of timing. Here, when examining the standard deviation of either the normalized position between short and long ratios, or the normalized force direction data, we observed a striking increase in variability for both groups throughout the presented interval that continuously increased throughout the interval. However, two notable differences between free-movement and hold groups were readily observed. First, whereas variability increased linearly in the hold group (Figure 7B), in the free-movement group this increase was slow to start, then rapidly increased before peaking at ^~^2800ms and then decreasing until the end of the 4000ms interval. Both increases are consistent with the increasing uncertainty in timing longer intervals, in accordance with Weber’s Law. Yet, the decrease in variability in the free-movement group is unexpected. One possibility is that, as the interval increases in length, free-movement subjects become very certain of it’s duration being longer than others in the stimulus set, and so variability of movement decreases as subjects approach the target. The second observation is that the momentary variability is far higher for the hold group than the free-movement group. This latter finding suggests that, if movement indexes timing, then increased variability of movement indicates greater uncertainty in the hold condition. This difference may further explain the difference in CV between groups; the allowance of free-movement improves temporal perception.

To further examine the predictability of movement dynamics for indicating timing and decision making, we again conducted a logistic regression analysis, but this time only for the 2000ms interval, which was located at the middle of the stimulus set and is thus the most ambiguous (Figure 8). Accordingly, movement dynamics should indicate which choice the subject will make. For the free movement group, distance ratios successfully determined which choice the subject would make up to 345ms prior to interval offset (all p < 0.05, Bonferonni corrected; 540ms uncorrected), demonstrating that by 1655ms or 82% of the way through the interval, the choice the subject would eventually make could be determined. For the hold group, force data revealed a different pattern; we were unable to decode subject choice at any point prior to offset of the interval, even at the point of offset itself. This finding suggests that, in the hold group, force data provides a less reliable index of the unfolding time estimation and decision making process. On the other hand, it alternatively suggests that subjects in the hold group were less certain of their decision while timing, matching the decreased precision of interval estimates in this group.

One additional reason for the lack of decoding accuracy in the hold group comes from the presence of change-of-mind trajectories in this group. If subjects changed their mind after the arm is released at interval offset, then the force direction data prior to offset will only be weakly informative to where subjects will eventually wind up. Yet, these changes-of-mind may also be detectable in the force response profiles in the form of bias. To investigate this further, we conducted our logistic regression analysis on force and movement profiles during the 2000ms period prior to interval onset of the middle stimulus interval. Here, we found no significant effects in the free movement group; however, we did detect significant differences in the hold group, specifically during the last 100ms prior to stimulus onset (all p<0.05, uncorrected). Examination of the force profiles during this time revealed a separation in the force direction between trials on which subjects would eventually choose “short” or “long”. Crucially, we observed that the force direction in this case was in the opposite direction of the response location subjects would eventually choose. That is, if subjects were eventually going to respond “long”, they were initially pointing more towards the “short” choice location. Although this effect was not strongly informative of choice, it suggests that the difference in force reflected biases in the initial pointing direction, from which subjects would eventually change their mind.

### Differences in starting location

As noted above, one explanation for the difference in precision between the hold and free-movement groups is that one group was allowed to be closer to the target locations. As such, the increase in threshold for the free-movement group may simply be the result of them feeling less pressure to make a speeded response, although no difference in RT was found for shorter durations between these groups. To address this possibility, we conducted an additional experiment (n=14) in which subjects performed two blocks of the hold condition, yet with different hold positions, keeping them near or far from the target locations (Figure 1C). If the difference in precision between groups was due to the proximity to the target, then subjects should be more precise in their estimates and have a correspondingly higher threshold when they are held closer to the targets. The results of this experiment demonstrated this was not the case, as no difference between the CV was detected between conditions [Wilcoxon Signed Ranks Test, *Z* = -0.31, *p* = 0.975, 10000 permutations]. No difference between BP values was detected (p>0.05), while a significant difference in RT was found, with subjects responding faster when the hold location was closer to the targets [*Z* = -3.296, p<0.001, Cohen’s *d* = 3.722]; we note that no interaction with duration was observed, here, contrary to the difference between free movement and hold groups in the previous experiments. Application of the DDM also found little influence of the hold location; importantly, no difference in the threshold parameter was detected [*Z* = -0.094, *p* = 0.925], whereas a significant effect was found on the non-decision time parameter [*Z* = -2.982, *p* = 0.001, Cohen’s *d* = 2.639]. No interactions with duration were found and no other parameters exhibited significant differences (all *p*>0.05). These findings suggest that the increase in precision for the free movement group was not due to the closer proximity these subjects were afforded to the response locations.

**Figure 9: Figure 9.**
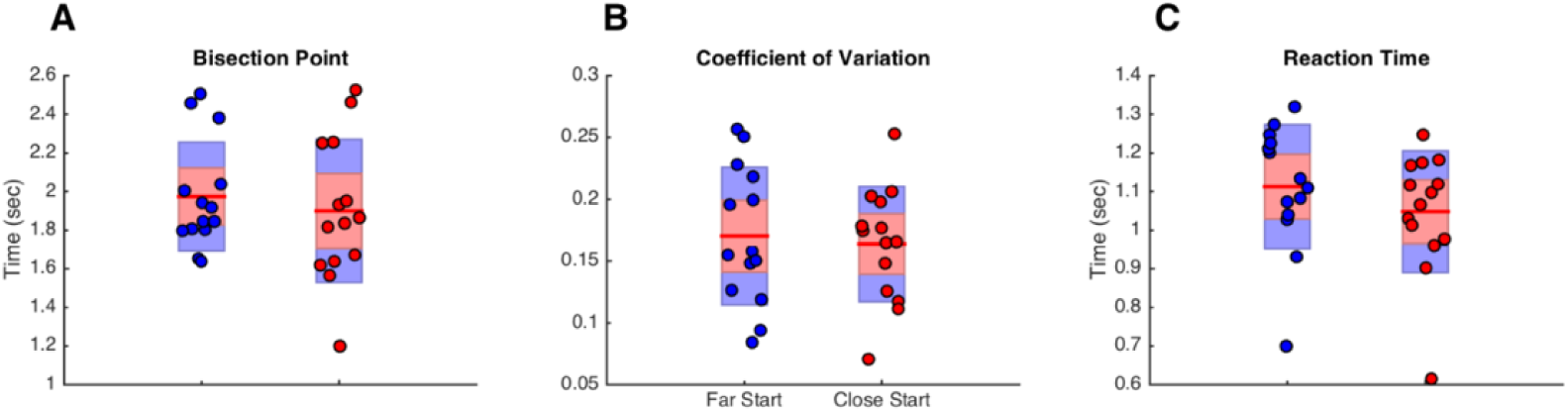
Comparison of behavior for different starting positions. Behavioral results from experiment 3 (different hold locations) are displayed for the **(A)** bisection point, **(B)** coefficient of variation, and **(C)** average reaction time across all durations. Red symbols represent individual subjects at the close hold location, blue symbols represent the long hold location. No significant difference between starting location was found for either the bisection point or coefficient of variation, but a significant difference was found for reaction time, with subjects responding sooner when the starting location was closer to the target locations.

## Discussion

Here, we show that when subjects are allowed to freely move while timing the duration of a concurrent auditory stimulus, they are more precise in their estimates than subjects who are held in place. We further link these findings to a change in the threshold parameter of a drift diffusion model (DDM) of temporal perception and decision making, with movement leading to a higher threshold for a response. These findings are not due to the closer proximity to the target or motor-based counting strategies in free-movement subjects. Further, we demonstrate that subject choice can be reliably determined by the statistics of movement dynamics prior to the offset of the interval, but only when subjects are freely allowed to move and not when held in place; further, movement variability increased as timed intervals grew longer, and was notably higher when being held in place. Lastly, we find that force exertion on the arm in subjects held in place can weakly predict eventual choice, but only in the period immediately before trial onset and in the opposite direction of the choice subjects will ultimately choose. These findings suggest that engagement of the motor system confers a perceptual advantage during time perception and decision making.

Previous work has begun to demonstrate a strong influence of the motor system on temporal processing. A variety of studies now demonstrate how movement parameters can influence the perceived duration of a stimulus, with longer movements typically associated with longer perceived durations (Yon et al., 2017; Yokosaka et al., 2015; Hagura et al., 2012). We find some support for this in our paradigm, with free-movement subjects classifying auditory tones as “long” more often when their arm movements during the interval were also longer. Yet, across subjects, those that moved more did not exhibit any difference in classification bias. One reason for this is that, in our task, subjects judged all intervals relative to a running average of previously presented ones; if subjects maintained consistent movement lengths throughout the task, then no difference across subjects would be observed. This raises the intriguing possibility that, if movement were perturbed during concurrent timing of an interval, the perception of that interval should change as well. Indeed, previous work using reaching paradigms have demonstrated that increased force resistance to a response location in a decision making task can lead subjects to choose that response less often, even when subjects don’t need to move the arm to make that choice (Hagura et al., 2017).

Beyond the effect of movement on altering the perception of time, our results here demonstrate an ability to improve it. Previous work by Hagura and colleagues (2012) has demonstrated that when subjects are preparing to make a movement, temporal fidelity for visual stimuli is improved. Yet, these authors did not analyze or report whether a change in precision also occurs, only a directional shift in the psychophysical function. Our findings demonstrate a general improvement in precision when subjects are allowed to move during the timing of an interval. In a separate vein, Tomassini and colleagues (Tomassini et al. (2015); (Tomassini et al., 2017) have shown an improvement in the perception of visual contrast during movement preparation and execution; however, this improvement varies sinusoidally (^~^4Hz) around the time of movement, suggesting a linkage with visual sampling mechanisms (Busch and VanRullen, 2010). Further work by Yon and colleagues (Yon et al., 2017) demonstrated that concurrent movements while timing a sub-second auditory stimulus, a closer analogue to the present task, also shifted the perception of duration, depending on the length, speed, or direction of the movement; however, no overall differences in precision were detected between different movement conditions. Notably, these authors did observe a change in precision in one experiment, with subjects exhibiting *better* precision when *not* moving compared to moving, opposite to what we observed here. Yon and colleagues explained this finding by suggesting that the dual task nature of the experiment, wherein movements were primed to a particular location while concurrently encoding the auditory stimulus duration, increased the difficulty of discrimination compared to a condition without movement. This finding is consistent with previous work demonstrating that dual-task interference leads to a shortening of perceived time, as well an increase in the variability of temporal estimates (Brown, 1997). In the present study, subjects were simply allowed to move throughout the interval, but no particular pattern or strategy was encouraged, with some subjects choosing not to move at all during the interval.

With respect to the potential neural mediators of the effects in our study, we note that recent evidence suggests that neurons within the supplementary motor area (SMA) code for subjective duration (Mita et al., 2009), and EEG activity putatively driven by the SMA has been shown to vary on a trial-by-trial basis with duration categorization (Wiener and Thompson, 2015). Further evidence has recently revealed that SMA neurons encode a categorical boundary for classifying intervals (Mendoza et al., 2018). Yet, alternative explanations suggest this region more broadly encodes response caution, and the evaluation of evidence in favor of a decision (van et al., 2011). In the motor control literature, the SMA has also been linked to sequential motor actions, and to an online “correction” of a motor plan following an error or perturbation (Nachev et al., 2008). Further evidence demonstrates that SMA neurons serve an action-monitoring role, in which they continuously evaluate the success of planned actions (Bonini et al., 2014). In the present study, the allowance of free-movement may have provided these subjects with the ability to evaluate choice in a continuous way. This possibility matches recent suggestions that the use of the motor system refines our perception of time (Coull et al., 2016). Evidence for this possibility comes from our finding that inter-trial variability of movements in the free-movement group during the evaluation of the interval were dramatically lower than in the hold group, suggesting a more precise representation of time for categorizing the interval. Similar work in animals has demonstrated that, when moving, the tuning of visual representations is enhanced (Dadarlat and Stryker, 2017). Indeed, a growing number of studies now demonstrate the enhancement of sensory representations during active behaviors across several species (Busse et al., 2017).

Based on the above findings, one explanation is that use of the motor system allowed free-movement subjects to “simulate” the timed interval, taking advantage of the greater temporal fidelity of the motor system. Yet, the timing of auditory stimuli alone is already highly precise, moreso than for visual (Wiener et al., 2014) or tactile (Jones et al., 2009) modalities. However, when combined, previous work has demonstrated an improvement over unisensory stimuli alone (Occelli et al., 2011; Manning and Schutz, 2013). Similarly, when tactile information is presented bimodally with visual stimuli, the perception of duration is also improved over unisensory conditions (Ball et al., 2017). Further, experience with audiomotor timing may confer additional advantages; professional drummers are notably less biased and more precise in the timing of both auditory and visual stimuli (Cicchini et al., 2012). Likewise, auditory frequency discrimination performance is impaired when subjects receive stimulation to the somatosensory cortex, but only when they are required to simultaneously attend to touch (Convento et al., 2018). In a similar vein, recent research has begun to demonstrate that time perception is highly linked to the amplitude and phase of beta (13-30Hz) oscillations. That beta oscillations are also highly implicated in movement is likely no coincidence; movement and timing may be intrinsically linked. Moreover, beta oscillations are invoked for the timing of durations even when no timed motor response is required, and stimulation at beta frequency (20Hz) shifts the memory of the perceived durations (Wiener et al., 2018a).

If the motor system is intrinsically involved in time perception, then why does the free-movement condition confer an advantage over the hold condition? That is, if timing is already partially instantiated in the motor system, then why should additional movement provide better temporal resolution? In the hold condition, the motor system may be computing parallel yet competing action plans for separate movements to the different target locations, as is the case for other variable reach targets (Cisek and Kalaska, 2010). In the present study, at the start of a trial, the reaching plan to the “short” target location should be dominant, but as the interval elapses the “long” target reach plan should gradually start winning, until the internal criterion is passed and subjects decide that the interval is now “long”; evidence for these competing action plans can be observed in the force direction data from the hold group, which shows a gradual transition, rather than a stepwise shift from short to long. Yet, in the free-movement group, this competition may not necessarily be taking place, as subjects are here able to fluidly explore the 2-dimensional space between targets while sequentially sampling information in favor of one versus the other; that is, in the hold condition, movement preparation precedes initiation, whereas initiation precedes preparation in the free-movement condition (Haith et al., 2016). In this case, noise arising from the competitive process may have less of an influence on the accumulation of evidence (Brunton et al., 2013).

One additional noteworthy difference between hold and free-movement subjects is the clear presence of change-of-mind trajectories in the hold condition. This observation does not preclude changes-of-mind in the free-movement condition, only that they are more apparent when subjects follow a predictable trajectory, matching the previous literature (Resulaj et al., 2009). Changes-of-mind in decision making have previously been suggested as a post-stimulus evaluation of evidence, in which sensory evidence “catches up” with the decision-making apparatus. However, time perception offers a special case for decision making, as subjects will experience a continuum of every interval in the stimulus set on a trial with the longest interval; as such, when a stimulus extinguishes, if subjects continue to accumulate time, they should only ever change from a “short” to a “long” choice, and not the other way around. In our study, changes-of-mind can be observed in both directions. Further, our single-trial analysis of the hold condition revealed that subject choice for the most ambiguous interval in our stimulus set (2000ms) could be decoded from directional force prior to interval onset, with subjects pushing in the opposite direction of the choice they would eventually make. This findings suggests that changes-of-mind occurring in the hold condition were not the result of a post-stimulus evaluation of evidence, but instead come from pre-stimulus biases in movement direction, perhaps from the previous trial (Wiener et al., 2014).

In the present study, the application of a drift-diffusion model of timing and decision making allowed us to dissociate to some degree the source of the differences between hold and free-movement conditions. Here, our model-based analysis suggests that, in the free-movement group, a higher threshold value is afforded, such that more evidence is required for subjects to commit to a response choice. Previous evidence suggests that increases in the response threshold translates to a reduction in impulsivity (Cavanagh et al., 2011); similarly, changes in the threshold parameter match the speed-accuracy tradeoff, with higher thresholds for decisions emphasizing accuracy (Voss et al., 2004). Yet, in the present study, accuracy was only relative to how well a subject could segregate around their own subjective internal criterion, and no feedback was ever provided. Further, free-movement subjects responded faster than hold subjects, but only for durations longer than the bisection point, whereas the increase in threshold was globally independent of duration. As such, while the drift-diffusion model findings suggest that free-movement subjects were more cautious in their response choices, we suggest that our findings cannot alone be explained by a change in decision strategy.

Despite heterogeneity in the free-movement strategies across subjects, we found that continuous proximity to the target location served as a good predictor of eventual choice. For the middle and most ambiguous interval in the stimulus set, choice location could be determined well before the stimulus offset occurred. Notably, longer movements on these trials more often led subjects to classify the stimulus as “long”, concordant with previous findings (Yokosaka et al., 2015; Yon et al., 2017). However, even when taking this into account, the divergence between target proximities is striking, with a separation by 1500ms into the trial that is predictive of choice. Given that subjects don’t know that the interval will end at 2000ms, this suggests that subjects commit to their choice based on proximity to the target at trial offset. As subjects adopted consistent movement strategies throughout the experiment, then the choice of where to respond becomes easier when interval offset occurs. This may in turn translate to a steeper categorization gradient and a smaller coefficient of variation, as observed for free-movement subjects.

In summary, we report here a dissociation in the precision of temporal estimates when subjects are allowed to freely move during the presentation interval compared to being held in place. Our findings cannot be explained by counting strategies or proximity to the target, and may depend on an adoption of the motor system to improve the fidelity and categorization of perceived time intervals. These results have implications for decision making and time perception studies, and suggest that the motor system is intrinsically linked to the perception of duration (Merchant and Yarrow, 2016) and that its adoption can aid decision making (Wolpert and Landy, 2012).

## Materials and Methods

### Participants

A total of 54 healthy subjects (27 male and 27 female, 23 ± 0.41 years of age) with no known neurological disorders or impairments were recruited for this study. All participants were right-hand dominant and used this hand to perform the task throughout the entirety of the experiment. Each participant only performed a single experimental paradigm and was naïve to the task. The study protocol was reviewed and approved by the George Mason University Institutional Review Board.

### Apparatus

Participants were seated in an adjustable chair directly in front of the robotic manipulandum (KINARM End-Point Lab, BKIN Technologies) at a height where their forehead could rest comfortably on the system’s headrest. A horizontal mirror display occluded the subject’s view of the right forearm to limit feedback of the upper limb and hand position to only what was observed on the screen. Visual feedback from the task was projected onto the mirror from a downward-facing LCD monitor positioned directly above. Participants gripped the right handle of the robotic manipulandum and made reaching movements to one of two circular targets 0.5 cm in diameter, placed 14 cm apart on the sagittal axis of the body (left target is the short duration location and right target is the long duration location, Figure 1C). During movements, the manipulandum continuously measured hand position, velocity, and forces applied by subjects while simultaneously exerting external forces at a sampling rate of 1000Hz.

### Experiment 1 - Free-Movement

In the first experiment (Figure 1A), termed the “free-movement” group, subjects (n=20) were first instructed on each trial to move the robotic manipulandum so that the screen cursor was brought to a starting location. Once there, the manipulandum was locked into place and subjects were unable to move the handle. Subjects remained locked in this position for 1000ms while the word “Get Ready” was presented 10 cm under the starting location. Next, the short and long duration locations were presented to the subject; both cues were equidistant from the starting position, located at 105° and 75°, respectively. After, the ready cue disappeared, the starting location turned blue and the robotic manipulandum was released from the hold position, allowing subjects to freely move. This “warm-up” period lasted for 2000ms, after which a tone was played via speakers to the subject. Tone durations were logarithmically spaced between 1 and 4 seconds in seven steps, and presented in a first-order counterbalanced sequence (Aguirre et al., 2011) for a total of 56 trials for each duration (392 trials total divided into 8 blocks of 49 trials each). Once the tone duration had elapsed, subjects were required to classify it into short or long duration categories. Subjects were instructed to judge each duration relative to the average of all previously presented durations. Notably, if a subject moved into either the short or long duration target locations during the warm-up or any other time before the to-be-presented duration had elapsed, the two duration locations disappeared and subjects were required to start the trial over again. All subjects were additionally instructed that they must respond as quickly, yet as accurately, as possible. As such, the ideal strategy is to move the cursor as close to the “short” target as possible, and then wait until the subject is certain that the duration is not “short” but “long”. Once subjects made a choice, the corresponding target turned green and then all stimuli were removed. A new trial was initiated after a 500 ms inter-trial-interval.

### Experiment 2 - Hold

In the second experiment (Figure 1B), termed the “hold” group, a different set of subjects (n=20) performed the same task as the free movement group above, except that no warm-up period was provided, and the cursor was not released from the hold position until the offset of the tone duration. Thus, unlike the warm-up period in the free-movement group, subjects were not allowed to move into a readied position to respond (i.e., enter the short or long duration location).

### Experiment 3 - Different Hold Locations

In the third experiment (Figure 1C), termed the “different hold locations” group, a different set of subjects (n=14) performed the same task as the hold group above, except that start position changed across blocks (close start: 7.8 cm away from the duration category targets, far start: 10 cm away from the targets). As in the hold experiment, the cursor was not released from the hold position until the offset of the tone duration. In this condition, subjects completed twice as many trials as in the previous experiments, with 392 trials for each of the starting locations (784 trials total).

### Analysis

#### Behavioral Data

Behavioral data were analyzed similarly to earlier work with this paradigm (Wiener et al., 2014). Choice and reaction time data were first calculated for each subject for each of the seven tested durations in our stimulus set; reaction time data were estimated as the difference between the offset of the presented interval and arrival at the target location. Psychometric and chronometric curves were then generated for each participant. Psychometric curves were generated by plotting the proportion of long response choices for each of the seven tested durations; these points were then fitted by a cumulative Gumbel distribution using the *psignifit* version 3.0 software package (see http://psignifit.sourceforge.net/) for Python (˙). The Gumbel distribution was chosen to reflect the log-spacing of the duration stimuli, as well as the uncertainty associated with longer stimulus durations. Upper and lower thresholds, the approximate points at which the subject is 25% or 75% likely to judge the stimulus as long, were calculated using the bias-corrected bootstrap method implemented by psignifit, based on 1999 simulations. The results of this analysis yielded the bisection point (BP; the time value at which subjects were equally likely to judge the stimulus as long or short), the difference limen (DL; the difference between the upper [75%] and lower [25%] threshold values divided in half), and the coefficient of variation (CV; DL/BP). The BP thus reflects the subjective midpoint of the range of tested durations, while the CV reflects the normalized variability of measurements. Chronometric curves were constructed by plotting the reaction time for each of the seven possible durations.

#### Movement Data

Movement data were extracted for all experimental conditions by calculating the X and Y positions of the manipulandum, as well as the force applied in each direction. Force direction data was calculated by the inverse tangent of X and Y force data using Matlab’s *atan2d* function. To calculate a reference point for the free-movement condition, we measured the Euclidean distance between the X/Y coordinate and the location of the short and long duration targets, and then took the ratio of the two (short/long). For movement length, we calculated the average Euclidean path length of movement during each of the seven intervals in our stimulus set.

As a measure of variability for both movement conditions, we calculated the average inter-trial standard deviation of movement location. This was done by first selecting data only for the last 100ms of each trial, and then calculating the standard deviation of the X and Y positions across trials for every 1ms timepoint leading up to arrival at the target location. We then calculated the average standard deviation across this 100ms interval for each of the seven stimulus intervals in our stimulus set.

#### Drift Diffusion Modeling

Drift diffusion modeling (DDM) was accomplished using routines provided by the HDDM package for Python (Wiecki et al., 2013). Model construction was conducted using the *HDDMStimCoding()* class, in which we created a model of choice and reaction time with duration as a within-subject feature. Specifically, we generated a model with four parameters: *a* (threshold), *v* (drift rate), *t* (non-decision time), and *z* (starting point), with each parameter varying by the tested duration. Our choice of this model conforms to earlier work on the DDM as applied to temporal bisection data (2ı); (Tipples, 2018), as well our own work demonstrating that this model provides the best fit to bisection data (Wiener et al., 2018a). Model fitting was applied individually to each subject across all three experiments; for experiment 3, the four parameters were additionally set to vary by the starting location (Far/Close). Fits for parameter values were minimized using *HDDM.Optimize()* for quantile optimization, which applies Maximium Likelihood estimation to the cumulative distribution of reaction times (Ratcliff and Tuerlinckx, 2002).

#### Statistical Analyses

For all bivariate comparisons between groups, we conducted two-tailed Mann-Whitney U-tests. For within-subject bivariate comparisons, two-tailed Wilcoxon Signed Rank Tests. In both cases, alpha was set at 0.05, and all *p* values were confirmed with 10,000 permutations to generate 95% confidence intervals. For comparisons of reaction time, where duration was a factor, mixed model ANOVAs were run with duration as a within-subject factor and group as a between-subject factor. For experiment 3 ( different hold locations), duration and distance were the only within-subject factors. Measures of effect size were calculated for all significant effects.

#### Logistic Regression Analysis

To conduct regression for choice data based on single-trial movement trajectories and force data, several steps were taken. First, a softmax transform was applied to the ratio of short/long distance values for the free-movement group, and the force direction data for the hold group, to constrain extreme values and bring both sets into a comparable reference frame. Next, we extracted these data in 1ms steps for the 1000ms prior to offset for each of the seven intervals in our stimulus set. This timeframe was chosen as it represents the minimum number of time points that can be viewed across all seven intervals, as the shortest interval in our stimulus set was 1000ms. Then, for each 1ms time point, we constructed a Generalized Linear Mixed Effects Model, using the *fitglme* function for Matlab. Initially, each model was constructed such that the choice on any trial (Y) could be predicted by a linear combination of the duration presented on that trial (D) and the normalized short/long ratio for free-movement and the normalized force direction for the hold group (M) as fixed effects, with individual subject (S) as a random effect, resulting in the function [Y ^~^ 1 + D + M + (1|S)] in Wilkinson notation. A binomial distribution was chosen to model the response distribution of choices, with a logit curve as the link function. Fitting of each model was accomplished using the Laplace approximation for Maximum Likelihood, as optimized by the Matlab function *fminsearch*. The result of this analysis yielded 1000 tests for each time point for free-movement and hold groups. To account for multiple comparisons, we applied a Bonferroni correction to maintain a testwise alpha level of 0.05, leading to an adjusted alpha of 0.00005. Following our initial analysis, we repeated our logistic regression analysis using only data from the 2000ms interval trials. In this case, the duration of the trial is always fixed, and so we calculated a reduced function without duration, using only the movement parameter for that subject. We again repeated this analysis using 1000ms. An additional analysis was run on the pre-onset time period for the hold group, which spanned 2000ms.

